# Tyrosine phosphoproteome profiling identifies cell-intrinsic signals limiting the efficacy of tyrosine kinase inhibitor therapies

**DOI:** 10.1101/2025.08.22.670548

**Authors:** Cameron T. Flower, Forest M. White

## Abstract

Tyrosine kinases (TKs) are frequently mutated or overexpressed in cancer, and TK inhibitors (TKIs) are an important therapeutic modality against TK-driven cancers, but many patients show an underwhelming response to TKIs prescribed on the basis of tumor genotype. To find cell-intrinsic TK signaling patterns which might be predictive of poor response to TKI exposure, we used high-sensitivity multiplexed mass spectrometry to quantify endogenous levels of 1,222 phosphotyrosine (pY) sites across the proteomes of TK-driven human cancer cell lines with variable response to genotype-matched TKIs. In direct comparisons between TKI-tolerant and TKI-sensitive lines with a common driver TK, we found that TKI treatment was equally effective at blocking driver TK signaling, and higher basal activity of the driver TK did not always predict higher sensitivity to TKI. All tolerant lines showed a dampened proteome-wide pY response to TKI exposure compared to sensitive lines, suggesting tumor cells with more robust TK signaling are less vulnerable to driver TK blockade. We found that each tolerant line depends on a unique set of compensatory TKs and signaling axes but are unified by hyperactivity of at least one of the SRC family kinases (SFKs) or the related ABL1/2 kinases, both at rest and under TKI treatment, despite absence of SFK/ABL genetic mutations. In time- and dose-resolved drug combination experiments, SFK/ABL inhibitors were potently synergistic with all TKIs tested, demonstrating that elevated SFK/ABL signaling is a conserved bottleneck for maximal TKI efficacy which could be exploited therapeutically.

**Significance:** The last twenty-five years have seen a remarkable number of new anti-cancer therapies specifically targeting tyrosine kinases, which frequently drive tumorigenesis when hyperactive. While these therapies have brought undeniable improvements in survival time and quality of life for a large fraction of patients, many still do not respond to inhibitors aimed at their presumed driver kinase. By profiling tumor cell-intrinsic signaling networks using phosphotyrosine immunoaffinity pulldown and high-sensitivity mass spectrometry, we directly compare the signaling networks of drug-tolerant and drug-sensitive cancer cell lines across several oncogenic driver contexts and find both unique and shared signals promoting therapy tolerance.

## Introduction

Recognition of the cancer-causing potential of tyrosine kinases (TKs), beginning with the seminal discovery of *v-Src* (1, 2) and underscored by the landmark identification of *BCR-ABL1* as a highly effective drug target in chronic myeloid leukemia (3, 4), has placed great emphasis on the role of TK signaling in cancer initiation and progression. As a result, dozens of TK inhibitors (TKIs) are now approved in the U.S. and globally for the clinical management of multiple solid and hematological malignancies, constituting the largest class of small-molecule therapeutics (5). While these drugs have afforded many patients longer survival and superior quality of life compared to the preceding standards of care, response rates for even the most advanced agents still fall far short of 100% (often ranging from 40-80%), and the biological determinants of poor initial response to TKI therapy remain poorly understood (6–11).

Tumor cell-intrinsic signaling networks play an important role in targeted therapy response, principally by dictating the activity of pathways downstream from, or redundant with, the intended drug target; for example, TKI treatment may fail to arrest the growth of tumor cells in which mitogenic signaling can be sustained by an alternate kinase (12). While progress has been made in cataloging the mechanisms tumor cells can deploy to dampen TKI efficacy, we have yet to uncover combination therapies which durably restrain tumor cell proliferation for most patients, and it remains unclear whether drug-tolerant tumors cells driven by distinct oncogenic TKs exhibit shared TK signaling dependencies, which may represent new therapeutic targets with broad efficacy. Here, using high-sensitivity untargeted mass spectrometry to profile the activity of TK signaling networks at proteome scale, we exploited the intrinsic differences in TKI sensitivity across TK-driven human cancer cell lines to examine the relationship between tyrosine phosphorylation patterns and response to genotype-matched TKI treatment.

## Results

### Drug response characterization of TK-driven cancer cell lines

To examine the role of TK signaling networks in mediating cellular sensitivity or tolerance to TKI treatment, we assembled a panel of established cancer cell lines harboring genetic alterations in clinically actionable TKs. We chose lines derived from patients with non-small cell lung cancer (NSCLC) due to the high frequency of TK alterations in NSCLC, the broad array of approved TKI therapies for NSCLC treatment, and the wide availability of established TK-driven NSCLC lines. Using the Cancer Dependency Map (13), we considered lines with known gain-of-function alterations in *MET* (focal amplification), *ALK* (*EML4-ALK* fusion), or *EGFR* (kinase domain point mutations) and with reported differences in sensitivity to genotype-matched TKIs. (Hereafter we adopt the acronym gmTKI to refer specifically to TKIs which are directed against the known oncogenic driver TK of a given cell line.) Our final panel consisted of six cancer cell lines, two for each driver TK: H1993 and EBC-1 (*MET*); H2228 and H3122 (*ALK*); and H1975 and HCC4006 (*EGFR*). Under dose-resolved challenge with six clinically approved TKIs, we verified that driver TK status is generally predictive of whether a cell line will show any response to gmTKI (Fig. 1A). In one case, H1975 required substantially higher doses of afatinib to achieve the effect observed in HCC4006, consistent with the reduced activity of afatinib against the *EGFR*^T790M^ allele present in H1975 (14), indicating that this cell line panel recapitulates known allele-specific TK targeting.

**Figure 1.**
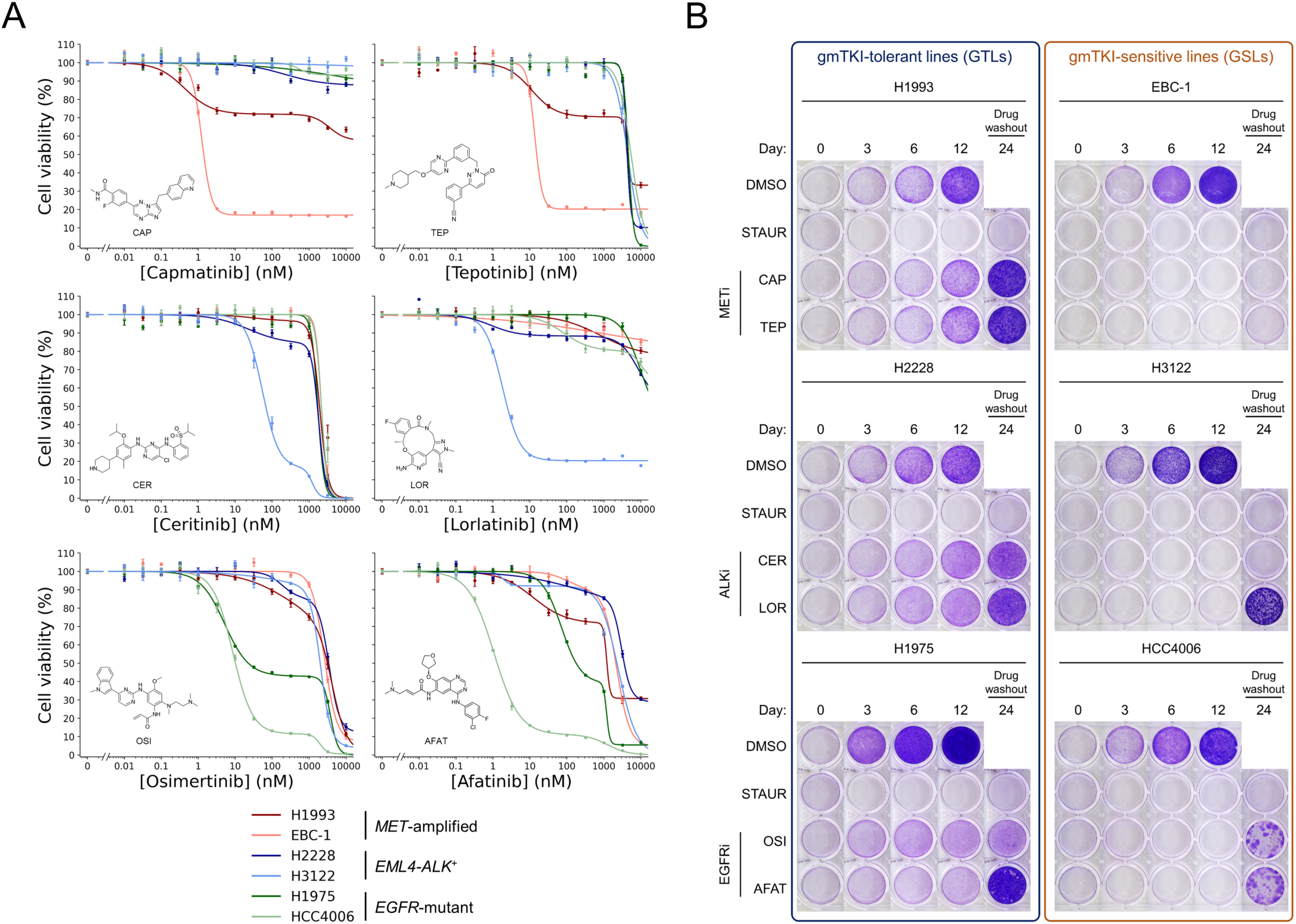
Tyrosine kinase (TK)-driven human cancer cell lines have substantial variability in TK inhibitor (TKI) sensitivity. (A) Dose-resolved cell viability measurements of cell lines under TKIs directed against (top) MET, (middle) ALK, and (bottom) EGFR. Treatment duration was 72 hours. Error bars depict standard error of the mean across six replicates. (B) Time-resolved clonogenic assays of TK-driven cell lines under genotype-matched TKI (gmTKI) exposure. All drugs were delivered at a concentration of 1 μM except staurosporine (10 μM).

Though all cell lines showed a response to gmTKI, within each pair of lines sharing a driver TK, one line appeared substantially more sensitive to drug than the other. Directly comparing dose-responses between cell lines is confounded by intrinsic differences in basal growth rate (15); we therefore used time-resolved clonogenic assays by crystal violet staining as an orthogonal qualitative approach, which confirmed that each pair of cell lines sharing a driver TK consisted of a gmTKI-sensitive line (GSL), which showed early and durable proliferative arrest or cell death from gmTKI exposure at a moderate dose (1 μM), and a gmTKI-tolerant line (GTL), which showed slow but sustained proliferation from the onset of exposure (Fig. 1B). A washout period following treatment confirmed that sustained gmTKI exposure is largely cytotoxic to GSLs, with some instances of subclonal drug tolerance resulting in colony formation from rare drug-tolerant persister cells (16). These results demonstrate that our TK-driven cancer cell line panel represents cell-intrinsic differences in sensitivity to various oncogenic kinase inhibitors.

### Co-occurring mutations contribute to gmTKI tolerance in a subset of GTLs

A common observation in tumor biopsies from patients who are unresponsive to targeted therapy is the presence of multiple co-occurring oncogenic mutations, which can result in distribution of proliferative and survival signaling across multiple pathways (17). Although co-occurring oncogenic mutations vary patient-to-patient, we wondered whether the GTLs in our panel harbored additional oncogenic mutations in signaling proteins which would offer a trivial explanation for tolerance to single-agent gmTKI. Using the Cancer Dependency Map, we found heterozygous point mutations in phosphoinositide 3-kinase (PI3K) family genes in two out of three GTLs: H1993 carries *PIK3CD*^E320D^, a mutation within the C2 domain of the catalytic subunit of PI3Kδ proximal to residues with known oncogenic potential when mutated, and H1975 harbors *PIK3CA*^G118D^, encoding a hyperactive form of the catalytic subunit of PI3Kα with oncogenic potential according to OncoKB (18). To test whether these mutations confer tolerance to gmTKIs, we asked whether co-inhibition of PI3K signaling boosts the efficacy of MET- or EGFR-directed TKIs in H1993 or H1975, respectively (Fig. S1A). In dose-resolved cell viability checkerboard assays, we detected no synergy in H1993 between gmTKI and inhibition of PI3Kδ, and moderate synergy in H1975 between gmTKI and inhibition of PI3Kα after three days of drug exposure (Fig. S1B). Under longer-term treatment at a fixed moderate dose, we found that co-treatment with gmTKI and PI3K inhibitor led to durable proliferative arrest specifically in PI3K-mutant lines (Fig. S1C). Treatment with PI3Kδ inhibitor alone led to growth arrest in all lines, likely due to pan-PI3K targeting at 1 uM; however, combined treatment of PI3Kα inhibitor and gmTKI led to substantial cell death only in PI3K-mutant lines. These observations together suggest that PI3K co-mutations explain some degree of gmTKI tolerance. Co-inhibition of the AKT serine/threonine kinases, which act downstream of active PI3K, phenocopied co-inhibition of PI3Kα, indicating a dependency on the PI3K-AKT axis in H1993 and H1975. Exposure of cells to the mTOR inhibitor rapamycin significantly augmented gmTKI activity in all three lines, and in some cases was effective as a single-agent, suggesting that all lines carry elevated dependency on mTOR activity for growth in the presence of driver TK blockade.

Our query of the Cancer Dependency Map also revealed that H2228 harbors biallelic deletion of *RB1*, a tumor suppressor with important regulatory roles in cell cycle progression and lineage fidelity, and loss of which has been linked to drug resistance (19, 20). Chemical and genetic screens have identified a synthetic lethal relationship between *RB1* and the serine/threonine Aurora kinases (AURKs) (21–23), which facilitate mitotic spindle assembly and chromosome segregation during mitosis. We therefore wondered whether a hyperdependency on AURK signaling might explain low sensitivity to driver TK inhibition in H2228 (Fig. S1D). Co-inhibition of ALK and AURKA by checkerboard assay showed synergy only at exceptionally high doses of AURKA inhibitor far greater than the reported IC_50_ (Fig. S1E) (24), and we detected no synergy with co-inhibition of AURKB. Longer-term exposure showed that AURK inhibition alone was broadly cytotoxic in all GTLs (Fig. S1F), suggesting that the tolerance of H2228 to gmTKI is not explained by compensatory AURK signaling.

### Quantification of cell-wide TK signaling under gmTKI by pY proteomics

Complete response to targeted therapy is dependent on successful drug-target engagement, durable inhibition of downstream signaling, and absence of compensatory mitogenic or pro-survival signaling. Direct analysis of TK cell signaling networks under therapeutic challenge can provide a readout of each of these crucial processes (25, 26). Given the near-binary difference in sensitivity to gmTKI within each pair of cell lines in our panel, we reasoned that a direct comparison of proteome-wide TK signaling between GTLs and GSLs would provide insight into the mechanisms underlying gmTKI tolerance, and a comparison across lines with different driver TKs might reveal the generality of those mechanisms. To quantify TK signaling networks, we exposed each line to a moderate dose of gmTKI or drug-free solvent (dimethyl sulfoxide, DMSO) for 24 hours, snap-froze and lysed cells, and subjected whole cell lysates to multiplexed proteomics sample preparation and two-stage enrichment for phosphotyrosine (pY), the endogenous product of TK enzymatic activity (Fig. 2A, see Materials and Methods). Phosphopeptides were analyzed by high-sensitivity untargeted mass spectrometry, and tandem mass tags enabled relative peptide quantitation across all samples within each plex. This approach generated three quantitative maps of proteome-wide pY abundance (Fig. 2B).

**Figure 2.**
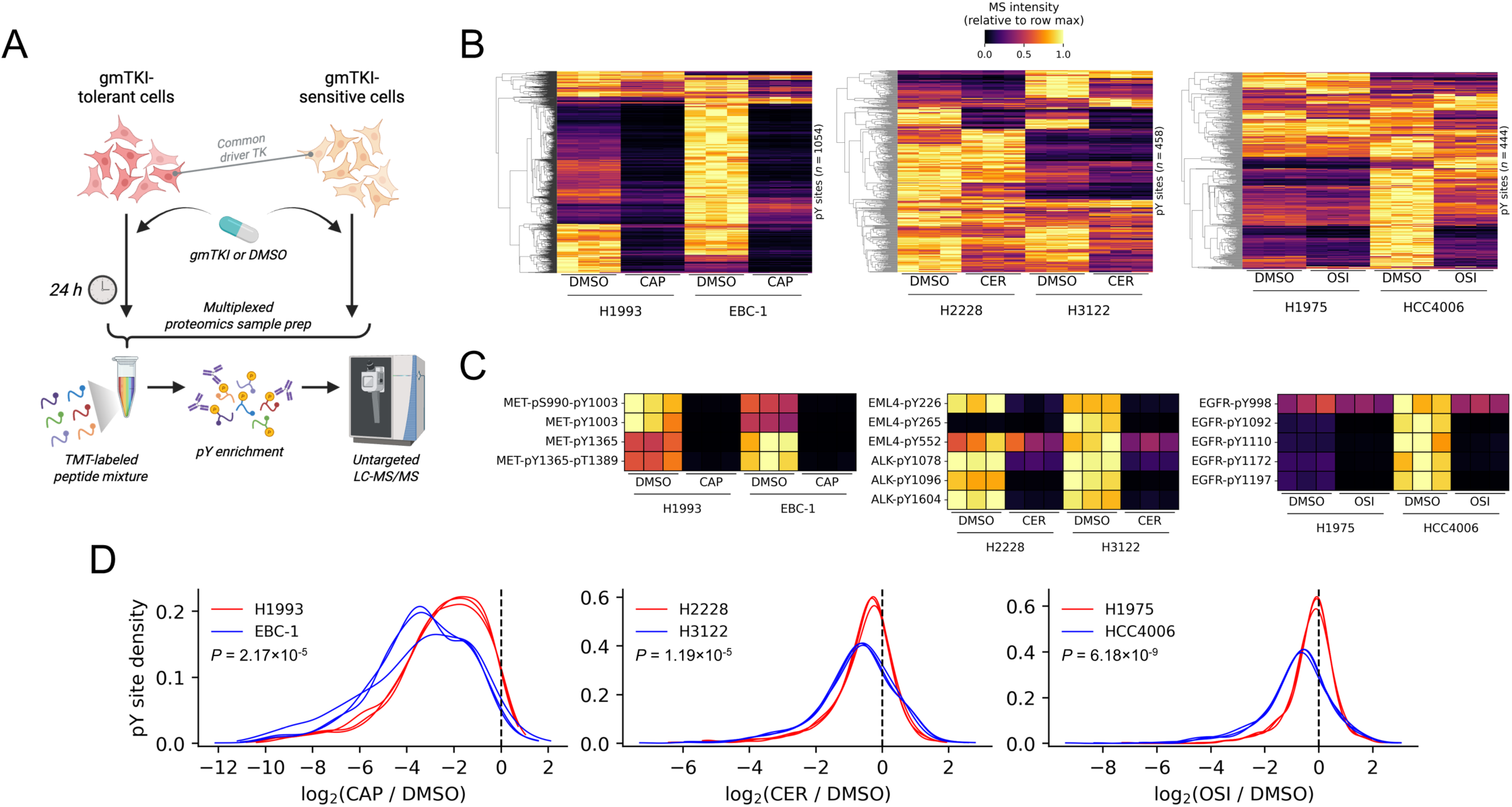
Tyrosine phosphoproteome profiling enables direct comparison of proteome-scale TK signaling patterns between GTLs and GSLs. (A) Experimental schematic depicting phosphotyrosine (pY) proteome profiling strategy by mass spectrometry. (B) Heatmaps depicting the relative abundance of all quantified pY sites, clustered by Euclidean distance, in cell lines driven by (left) MET, (middle) ALK, and (right) EGFR. (C) Heatmaps depicting the relative abundance of pY sites on driver TKs in cells under control solvent (dimethyl sulfoxide, DMSO) or gmTKI exposure. (D) Kernel density plots depicting the relative abundance distributions of pY sites in cell lines driven by (left) MET, (middle) ALK, and (right) EGFR under gmTKI compared to DMSO. *P*-values were derived by two-sided *t*-test.

Exploring each pY proteome, we first examined tyrosine phosphorylation directly occurring on each driver TK as a reporter of driver TK activity, in order to determine whether the driver TK was more completely inhibited in lines with the greatest sensitivity and whether GSLs owe their greater drug sensitivity to intrinsically higher driver TK activity. We found that in MET- and EGFR-driven lines, GSLs showed higher basal levels of driver TK autophosphorylation compared to GTLs, suggesting the possibility of elevated cell-intrinsic target activity, but this trend was not seen in ALK-driven lines (Fig. 2C). Under gmTKI treatment, we observed complete target inhibition in all lines, and a similar effect on phosphorylation of downstream adapter proteins and other signaling effectors (Fig. S2), demonstrating that tolerance in this system is not due to reduced drug import or ineffective target engagement.

While no significant differences in driver TK activity were detected between GTLs and GSLs under gmTKI, there was significantly more pronounced gmTKI-induced remodeling of the pY proteome in GSLs compared to GTLs (Fig. 2D). In all lines, most pY sites were downregulated by gmTKI exposure as expected, but cell-wide pY levels in all GTLs showed a greater robustness to driver TK inhibition, suggesting a general association between TK signaling fragility and vulnerability to gmTKI.

### pY proteome profiling enables detection of cell line-specific signals promoting drug tolerance

Among TK signaling events which were associated with gmTKI tolerance, we found that EGFR autophosphorylation – while significantly reduced by gmTKI in both MET-amplified lines, consistent with known MET-EGFR crosstalk (27, 28) – was residual in H1993 but not EBC-1, possibly due to greater basal EGFR activity in H1993 (Fig. 3A). Combining gmTKI with EGFR inhibition was synergistic (Fig. 3B) and led to durable growth arrest (Fig. 3C), indicating a high dependency on residual proliferative signal emanating from EGFR in the absence of oncogenic MET activity. These results agree with our observation of a partial response of H1993 to afatinib, which inhibits wild-type EGFR (Fig. 1A). In H2228, we detected sustained activity of two alternate receptor TKs, AXL and EGFR, under gmTKI (Fig. 3D). While inhibition of either one of these targets did not meaningfully augment inhibition of ALK, simultaneous inhibition of all three TKs led to a substantial reduction in growth rate (Fig. 3E), indicating that high basal AXL and EGFR activity contributes to gmTKI tolerance in this line.

**Figure 3.**
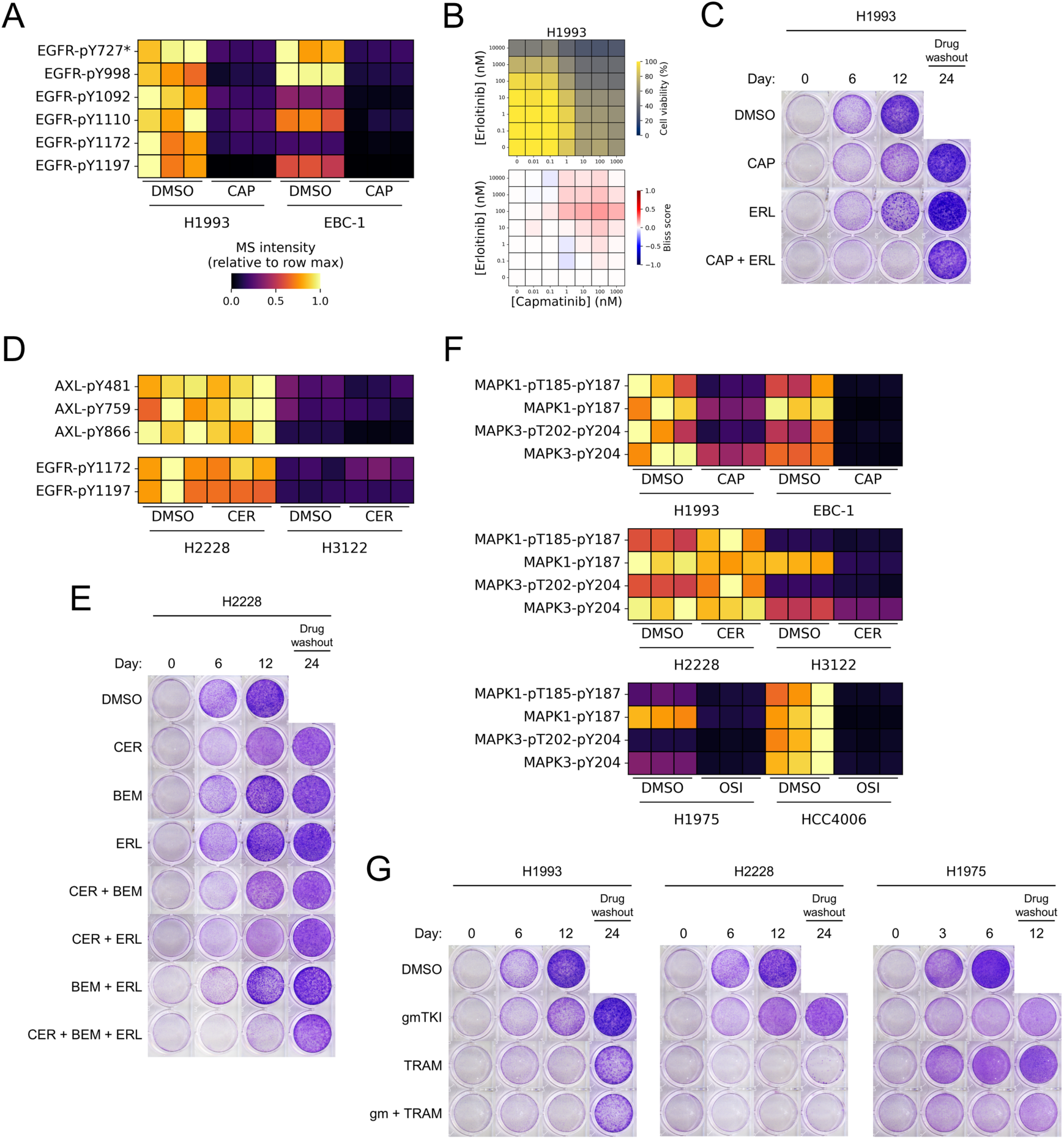
Tyrosine phosphoproteomics resolves signaling dependencies and effective anti-cancer drug combinations in GTLs. (A) Heatmap depicting the relative abundance of pY sites on EGFR in MET-amplified cell lines. Asterisk denotes a non-unique phosphopeptide mapping (EGFR-pY727; ERBB2-pY735; ERBB4-pY733). (B) Dose-resolved cell viability measurements (top) and corresponding Bliss synergy matrices (bottom) from checkerboard assays pairing gmTKI (capmatinib) with erlotinib in H1993 cells. Treatment duration was 72 hours. Cell viability values were averaged over six biological replicates and normalized to DMSO control. (C) Time-resolved clonogenic assay of H1993 cells treated with the gmTKI capmatinib, EGFR inhibitor erlotinib, or the combination. All drugs were delivered at a dose of 1 μM. (D) Heatmaps depicting the relative abundance of pY sites on AXL and EGFR in ALK-rearranged cell lines. (E) Time-resolved clonogenic assay of H2228 cells treated with the gmTKI ceritinib, AXL inhibitor bemcentinib, erlotinib, or combinations thereof. All drugs were delivered at a dose of 1 μM. (F) Heatmaps depicting the relative abundance of activation loop phosphorylation sites on MAPK1 (ERK2) and MAPK3 (ERK1) in all three GTLs. (F) Time-resolved clonogenic assay of GTLs treated with gmTKI, MEK1/2 inhibitor trametinib, or the combination. All drugs were delivered at a dose of 1 μM.

The RAS-ERK pathway is an important cascade that transduces signals from MET, ALK, EGFR, and many other receptors and cytoplasmic signaling pathways to coordinate cell proliferation, differentiation, migration, and other important cellular processes (29). High levels of phosphorylation on the activation loops of MAPK1 (ERK2) and MAPK3 (ERK1), canonically by the dual specificity kinases MEK1/2, are reliable indicators of RAS-ERK pathway activity. In H1993 and H2228, we observed ERK phosphorylation patterns which closely resembled EGFR and AXL phosphorylation (Fig. 3F), consistent with the role of RAS-ERK downstream of these receptor TKs. In H1975, where gmTKI alone led to complete inhibition of MEK1/2, addition of a MEK1/2 inhibitor expectedly failed to augment gmTKI; in contrast, MEK1/2 inhibition led to durable proliferative arrest in H1993 even without gmTKI, phenocopying the effect of dual MET/EGFR inhibition, and treatment of H2228 was potently cytotoxic. These results suggest a strong dependency on the RAS-ERK pathway in these lines (Fig. 3G).

### Basal and sustained SFK/ABL signaling is a unified driver of drug tolerance

Tumor-specific mechanisms of drug tolerance can be informative for considering tailored drug combinations but do not represent generalizable therapeutic strategies. To search for consensus signaling events promoting drug tolerance, we examined pY sites showing significantly greater abundance in all GTLs compared to the corresponding GSLs (Fig. 4A). Intuitively, several pY events known to drive cell cycle progression were higher in all GTLs under gmTKI compared to GSLs, including N-terminal phosphorylation of CDK1/2/3 (30) and activation loop phosphorylation of MAPK1/3 (31).

**Figure 4.**
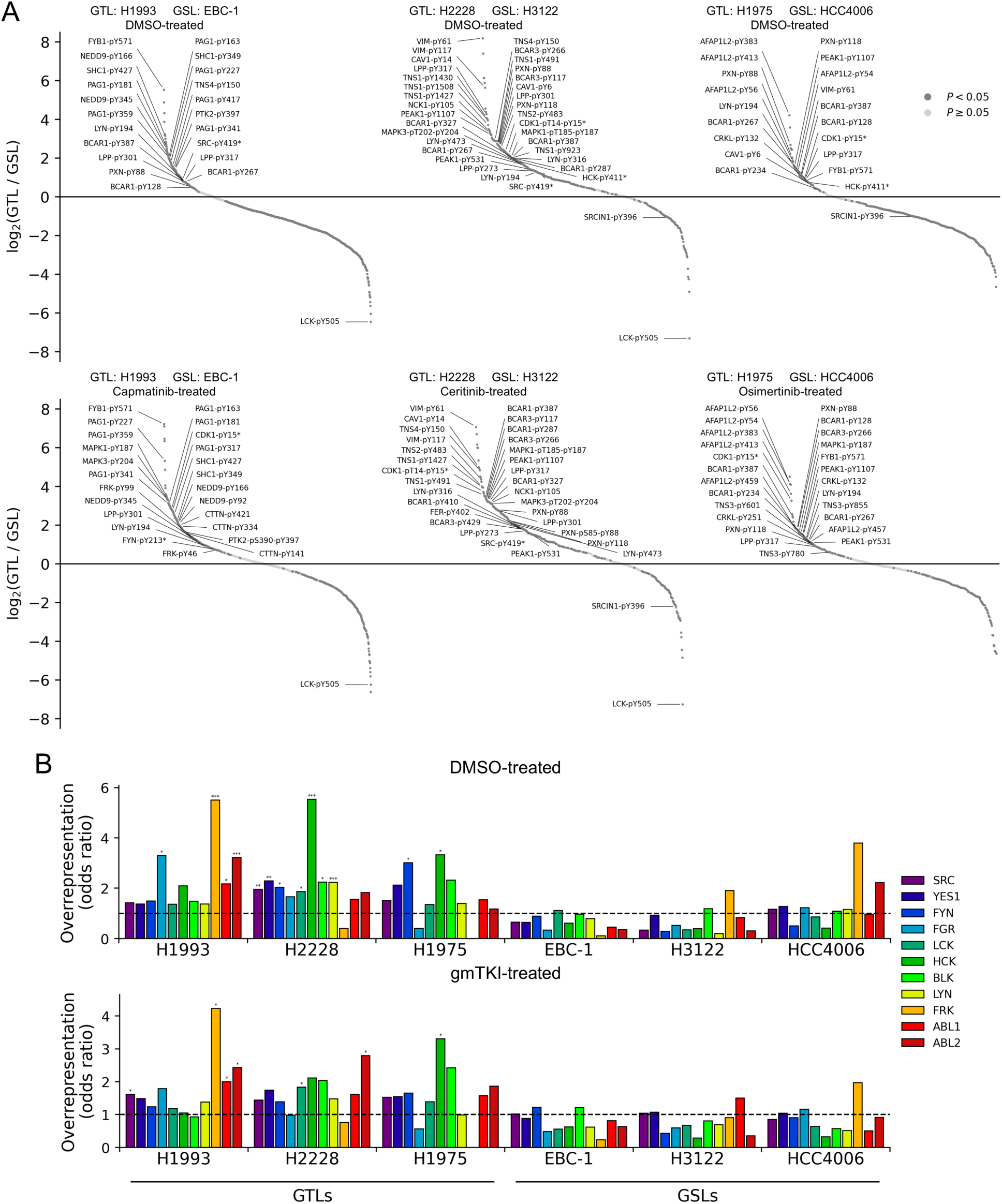
Elevated SFK/ABL signaling is feature of GTLs. (A) Scatter plots depicting pY sites rank-ordered by decreasing abundance in each GTL relative to the corresponding GSL, under exposure to (top) DMSO or (bottom) gmTKI. *P*-values were derived by two-sided *t*-test with multiple hypothesis correction by the Benjamini-Hochberg procedure. Asterisks denote non-unique phosphopeptide mappings. (B) Overrepresentation analysis quantifying the overlap between proteins with significantly elevated pY levels and those reported to interact with SFK/ABL kinases. Non-unique phosphopeptides were excluded from the analysis. \**P* < 0.05, \*\**P* < 0.01, \*\*\**P* < 0.005 by one-sided Fisher’s exact test with multiple hypothesis correction by the Benjamini-Hochberg procedure.

Many of the pY sites with elevated abundance in GTLs were on known substrates of the mitogenic and survival-promoting SRC family kinases (SFKs), including sites on BCAR1, LPP, NEDD9, and PXN, among others (32–35). Activating sites on SFKs, such as SRC-pY420, LYN-pY194, and HCK-pY411, also showed greater abundance in GTLs, and several SFK-inhibitory signals showed reduced abundance, together suggesting that heightened SFK signaling is a common feature of gmTKI-tolerant cells. We note that none of the cell lines in our panel harbored mutations in any of the SFKs according to the Cancer Dependency Map.

Since SFKs are known to phosphorylate hundreds of substrate proteins, we sought to test whether there was enrichment for SFK signaling proteins in GTLs compared to GSLs. Testing for enrichment of phosphorylated substrates of a given kinase remains challenging because most kinase-substrate relationships have yet to be discovered and reported at phosphosite-specific resolution (36); therefore we leveraged published protein-protein interaction data from BioGRID (37) to test for overrepresentation of known SFK interactors among pY sites which showed significantly greater abundance in GTLs compared to GSLs, since physical interaction between a kinase and its substrate is required for substrate phosphorylation. We included interactors of the SRC-like kinases ABL1/2, which are known to share substrates with the SRC family and regulate many of the same downstream processes (38, 39). We observed significant overrepresentation of SFK/ABL interactors in every GTL under DMSO and no significant overrepresentation in any GSL (Fig. 4B). Under gmTKI exposure, there remained at least one SFK/ABL kinase with significant enrichment within each GTL, and none in any of the GSLs. These results provide strong statistical support to the observation that high SFK/ABL signaling is associated with gmTKI tolerance.

Motivated by the result that basal and sustained SFK/ABL signaling is a common feature among GTLs, we paired each gmTKI with one of four SFK/ABL inhibitors in checkerboard assay format for each GTL. SFK/ABL inhibition was synergistic with gmTKI across all lines, demonstrating that elevated SFK/ABL signaling contributes to gmTKI tolerance (Fig. 5, A-D). The most synergistic combinations in H2228 and H1975 led to near-complete cell death. Under longer-term exposure at a moderate dose, we found that co-treatment with SFK/ABL inhibitors substantially augmented gmTKI activity in all GTLs, leading to durable cell cycle arrest or cell death in most cases (Fig. 5E). Together, these findings indicate that heightened basal SFK/ABL signaling, which is sustained under gmTKI exposure and not driven by SFK/ABL gene mutations, is a general and exploitable mechanism underlying drug tolerance.

**Figure 5.**
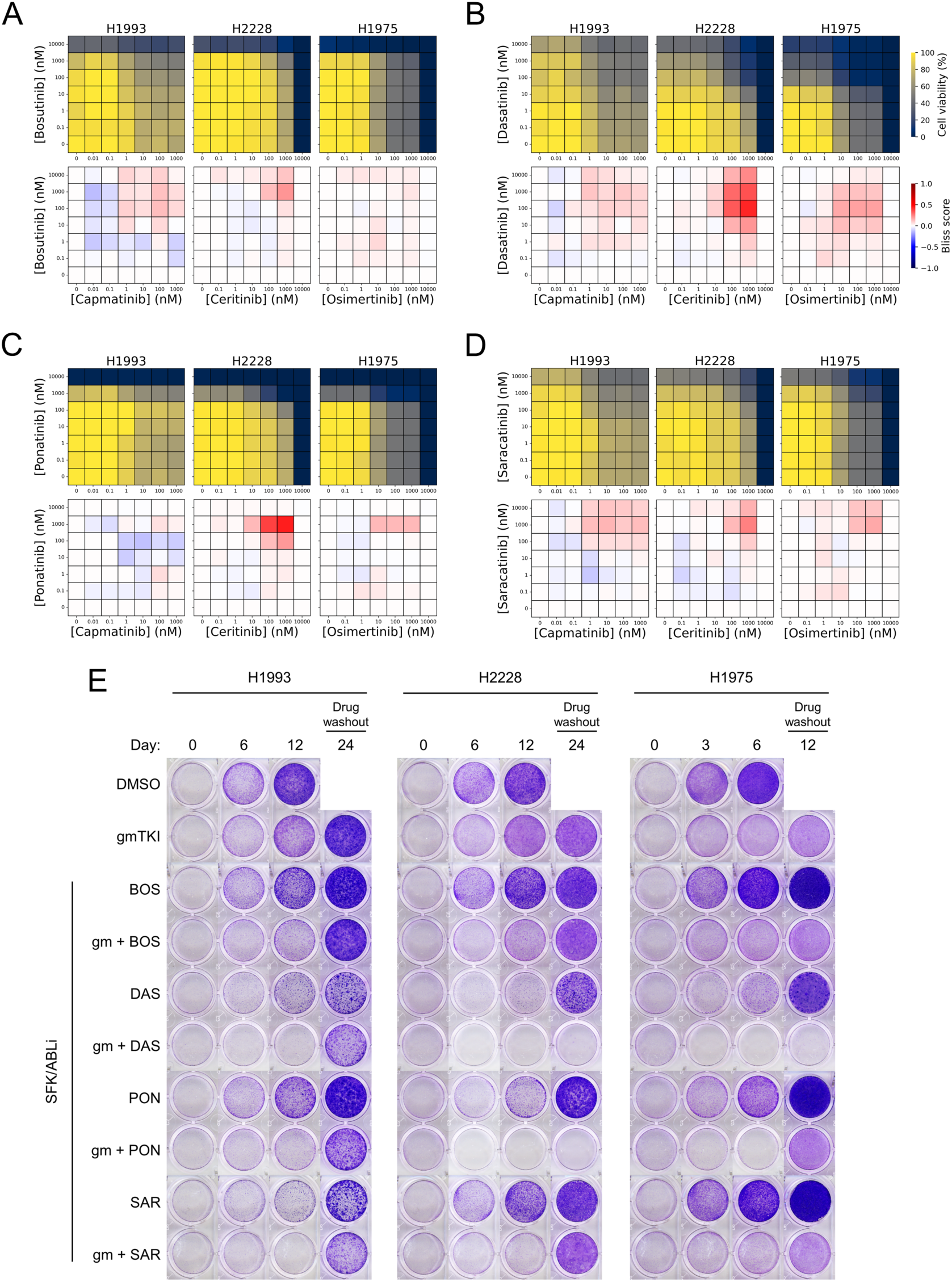
Pharmacological co-inhibition indicates that SFK/ABL signaling is a driver of gmTKI tolerance. (A-D) Dose-resolved cell viability measurements (top) and corresponding Bliss synergy matrices (bottom) from checkerboard assays pairing gmTKI with the SFK/ABL inhibitors (A) bosutinib, (B) dasatinib, (C) ponatinib, and (D) saracatinib in all three GTLs. Treatment duration was 72 hours. Cell viability values were averaged over six biological replicates and normalized to DMSO control. (E) Time-resolved clonogenic assays of GTLs treated with gmTKI, SFK/ABL inhibitors, or combinations thereof. All drugs were delivered at a dose of 1 μM. Timing of the experiment was compressed for H1975 due to its significantly faster growth which caused poor cell adhesion under certain drug treatments after 10-12 days.

Finally, we wondered about the relative contribution of ABL1/2 to the efficacy of SFK/ABL inhibitors combined with gmTKI. To isolate the effects of ABL1/2 inhibition from SFK inhibition, we treated cells in checkerboard assay format with imatinib, which is highly selective for ABL1/2 over SFKs, and found modest synergy with gmTKI in H2228 (Fig. S3A), consistent with heightened ABL2 signaling in H2228 under gmTKI (Fig. 4B). Under sustained exposure, imatinib co-treatment led to reduced growth uniquely in H1993 (Fig. S3B), consistent with heightened ABL1/2 signaling in this line (Fig. 4B). Altogether, these results demonstrate that elevated cell-intrinsic activity of SFK/ABL, likely in concert with other biological processes, represents a generalizable mechanism by which drug-tolerant cancer cells survive and maintain proliferative activity under the stress of TKI therapy.

## Discussion

In this work, we used quantitative mass spectrometry and exploited intrinsic drug sensitivity differences between human cancer cell lines to detect tumor cell-intrinsic signals contributing to targeted therapy tolerance. We focused on tolerance to TKIs, currently the largest class of small-molecule targeted therapy in the oncology clinic, and through direct measurement of TK activity by quantifying tyrosine-phosphorylated peptides across the proteome, we report signals which explicitly relate to this class of therapy. Our findings address several gaps in our current understanding of TKI tolerance. First, the molecular and cellular basis for acquired TKI resistance – that is, the loss of TKI sensitivity over the course of treatment – has been thoroughly explored in recent decades (40–42); by comparison, less emphasis has been placed on understanding the failure of therapy to restrain proliferation of tumor cell populations from the onset of drug exposure. Second, many of the reported mutations and gene regulatory programs underlying TKI tolerance vary substantially across patients and depend on the particular oncogenic driver TK, as well as the particular TKI chemistry, stressing the importance of finding general survival strategies employed by tumor cells (42). Third, relatively little is known about tolerance to inhibitors of MET or ALK compared to EGFR, likely due to the relatively recent clinical approval of inhibitors directed against these two targets and their lower mutation frequency in cancer compared to EGFR (7, 10, 43).

Oncogenic co-mutations are a widely accepted model for tolerance to targeted therapy directed against a driver TK; by sustaining signaling independently of the driver TK, or otherwise dampening the dependency of the cell on the driver TK for cycling or survival, co-mutated oncogenic events pose a significant barrier to therapeutic success in the clinic (17). We show that co-mutations can contribute to gmTKI tolerance in a cell line-specific manner, and that in some cases – such as H2228, which harbors RB1 deficiency – co-targeting the alternate oncogenic event is not necessarily productive or possible. According to the Cancer Dependency Map, some of the lines we examined harbor other co-mutations we did not pursue, including loss-of-function mutations in *STK11* (in H1993) and in *CDKN2A* (in H1975), both of which are tumor suppressors and could in principle contribute to gmTKI tolerance.

A central finding we report here is that SFK/ABL signaling is a shared dependency in GTLs, both with and without gmTKI exposure. The experiments demonstrating this result were motivated by the observation that each GTL showed high levels of phosphorylation of putative SFK/ABL substrates compared to the paired GSL. We note that GTLs showed phosphorylation of overlapping but unique sets of SFK/ABL substrates, likely a consequence of cell line-specific SFK/ABL expression, activity, subcellular localization, or substrate expression (44, 45).

Co-treatment experiments pairing gmTKI with SFK/ABL inhibitors showed that dasatinib and ponatinib effectively prevented outgrowth in all three GTLs, and drove complete cell death in a subset of lines. Bosutinib, however, did not show the same degree of efficacy despite being moderately synergistic with multiple gmTKIs. We reason this may be due to differences in target profiles across these agents; in particular, at least one SFK (LCK) is a confirmed target of dasatinib and ponatinib but not bosutinib according to DrugBank (46), and the detection of LCK-specific phosphopeptides in the pY proteome confirmed that each cell line tested here expresses LCK. Our results are therefore consistent with the expectation that different targeted agents have distinct polypharmacology.

Dasatinib and other SFK/ABL inhibitors have faced challenges in the treatment of solid tumors, principally due to on-target but off-tumor activity leading to dose-limiting toxicity (47, 48). Our findings suggest that there may be a therapeutic window for SFK/ABL inhibitors in patients with poor response to first-line TKI therapy, and a clinical assay to quantify SFK/ABL activity from tumor biopsies may be useful for patient stratification. Engineering tumor-specific delivery, for example by antibody-drug conjugation, could offer opportunities for reduced toxicity compared to systemic delivery.

One limitation of this study is that cancer cell lines, while easy to perturb and profile, are unable to recapitulate tumor cell-extrinsic signals emanating from the stroma, the host immune system, and other components of the tumor microenvironment. Recognizing this limitation, we chose to focus on cell-intrinsic signals, and we note that many of our observations are corroborated by clinical evidence, including reduced response rates to ALK inhibitors among NSCLC patients expressing EML4-ALK variant 3 (present in H2228) compared to patients expressing variant 1 (present in H3122) (49–51). Crucially, experiments performed using more complex *in vivo* model systems, as well as observations from patient biopsies, support our central finding of the importance of SFK/ABL signaling in shaping targeted therapy response (25, 52–59).

## Materials and methods

### Cell lines and drug treatment

H1993, H2228, H1975, and HCC4006 cell lines were purchased from ATCC. EBC-1 cells were purchased from JCRB Cell Bank. H3122 cells were provided by the Pasi Jänne lab (Dana-Farber Cancer Institute, Boston, MA). Cell lines were selected on the basis of their *MET*, *ALK*, and *EGFR* mutation status according to the Cancer Dependency Map (13). All cell lines were cultured in RPMI-1640 supplemented with 10% FBS and were regularly tested for mycoplasma contamination. All drugs used in this study were purchased from Selleck Chemicals and were dissolved in dimethyl sulfoxide (DMSO). In all drug treatment experiments, solvent exposure was held constant at 0.1% (v/v), and control cells were treated with 0.1% DMSO accordingly.

### Cell viability dose-response assay

Cells were seeded in a 384-well plate (500 cells/well, 3 drugs/plate, *n* = 6 replicates per drug dose and *n* = 18 replicates for control cells) and incubated for 24 hours, followed by treatment with drug at the designated doses for 72 hours. Cell viability was quantified using CellTiter-Glo (Promega, catalog # G7572) according to the manufacturer protocol. All luminescence values were normalized to the average of the drug-free control wells. Outlier values were removed according to the 1.5×IQR rule. Monophasic or biphasic dose-response curves were fitted to the data according to the following viability equations:

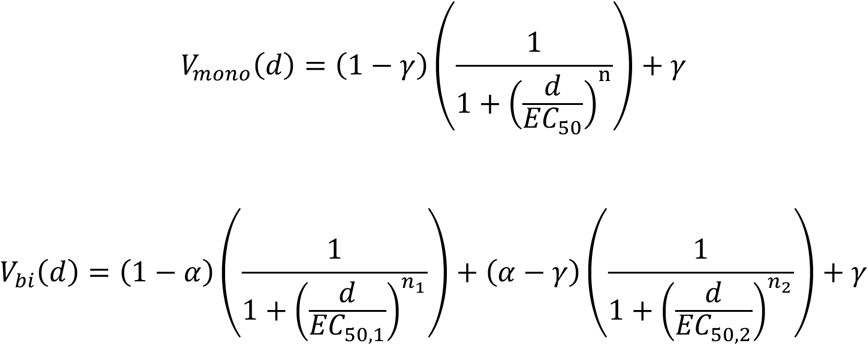

where *V*(*d*) ∈ [0,1] is the fraction of viable cells under dose *d* (in nM). Under the monophasic model, *EC*_50_ provides an estimate of the dose at which there is a half-maximal effect, and *n* is a measure of the steepness of the response. Under the biphasic model, *EC*_50,1_ and *EC*_50,2_ are estimates of the doses at which there is a half-maximal effect for the first and second dose-response phases, respectively. *α* provides an estimate of the intermediate viability of cells treated with doses between *EC*_50,1_ and *EC*_50,2_. *n*_1_ and *n*_2_ are measures of the steepness of the first and second dose-response phases, respectively. Under both models, *γ* provides an estimate of the minimum viability. Model fitting was performed using the curve_fit function in the SciPy scientific computing python package (60).

### Clonogenic assays

Cells were seeded in 24-well plates (20,000 cells/well) and incubated for 24 hours prior to drug treatment. Staurosporine (STAUR), a naturally occurring broad-spectrum kinase inhibitor, was included as a positive control for cytotoxicity. Drugged media was replenished every three days until drug washout, at which point all drugged media was removed and the cells were rinsed and replenished with drug-free media, which was replenished every three days. At each indicated timepoint, plates were removed from culture, media was removed, and cells were immediately fixed with 50% methanol in double-distilled water for 10 minutes. Cells were then stained with 0.5% crystal violet in 25% methanol for 30 minutes. Because many of the clonogenic assays were performed simultaneously, several of the image segments of DMSO- and gmTKI-treated cells are repeated in Figures 3, 5, S1, and S3. All raw images will be provided as supplementary material.

### Checkerboard assays

Cells were seeded in a 384-well plate (500 cells/well, *n* = 6 replicates per dose) and incubated for 24 hours, followed by treatment with drug pairs at the designated doses for 72 hours. The maximum dose used was 10 μM for all drugs except capmatinib (1 μM) due to its reduced solubility. Cell viability was quantified using CellTiter-Glo (Promega, catalog # G7572) according to the manufacturer protocol. All luminescence values were normalized to the average of the drug-free control wells. Outlier values were removed according to the 1.5×IQR rule.

For assessing drug synergy and antagonism, the Bliss independence model was used, which assumes that independent drug-drug interactions are multiplicative (61). The model is summarized as follows: if exposure of cells to drug A at a given dose results in effect *e_A_* ∈ [0,1], and exposure to drug B at a given dose results in effect *e_B_* ∈ [0,1], then under the null hypothesis that A and B act independently, the expected effect of combining A and B is given by

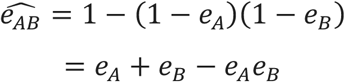

A and B are said to be synergistic if the observed effect *e*_,-_ of combining A and B (at their respective doses) is greater than the expected effect 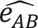 and antagonistic if the observed effect is less than 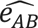. The degree of synergy or antagonism is therefore given by the Bliss score:

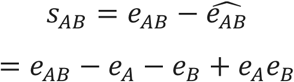

In experiments where relative cell viability at a fixed endpoint is measured or inferred, drug effect *e* at a fixed dose is related to viability *v* ∈ [0,1] as *e* = 1 − *v*, allowing the Bliss score to be directly expressed in terms of the normalized assay readout:

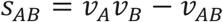

For each drug pair, Bliss scores were calculated and reported for all dose combinations contained within each 7×7 dose matrix.

### Phosphoproteomics sample preparation

Cells were lysed in 8M urea and total protein was quantified by bicinchoninic acid assay (Pierce, catalog # 23225). For each sample, 1 mg of protein was subjected to reduction in 10 mM dithiothreitol at 56°C for one hour, followed by cysteine alkylation in 55 mM iodoacetamide at room temperature for one hour in the dark. Samples were diluted 5X in 100 mM ammonium acetate to decrease the urea concentration prior to protein digestion, then samples were incubated with sequencing-grade modified trypsin (Promega, catalog # V5113) for 18 hours at a trypsin:protein ratio of 1:50 (w/w). Digestion was halted with 10% acetic acid, and samples were desalted by solid phase extraction using Sep-Pak C18 Plus Light cartridges (Waters, catalog # WAT023501) and eluted in 40% acetonitrile. All samples were concentrated in a vacuum centrifuge. Peptide concentration was quantified by bicinchoninic acid assay (Pierce), and 190 μg of peptide from each sample was freeze-dried by lyophilization. Samples were resuspended in 50 mM HEPES and labeled with TMTpro isobaric mass tags (Thermo Fisher Scientific, catalog # A52045) at a TMT:peptide ratio of 2.5:1 (w/w) for five hours at room temperature. Unreacted TMT was quenched with 3.2 μL of 5% hydroxylamine, then samples within each plex were pooled and dried by vacuum centrifugation. TMT channel assignment was randomized across samples within each plex.

### Two-stage enrichment of tyrosine-phosphorylated peptides

TMT-labeled samples were resuspended in immunoprecipitation (IP) buffer (100 mM Tris-HCl, 1% NP-40, pH 7.4). For IP of tyrosine phosphorylated peptides, samples were incubated with 24 μg of anti-pY 4G10 antibody (BioXCell, catalog # BE0194) and 12 μg of anti-pY PT-66 antibody (Sigma-Aldrich, catalog # P3300) conjugated to protein G agarose beads (Sigma-Aldrich, catalog # IP04). Immunoprecipitation was carried out over 18-24 hours at 4°C. Peptides were eluted with 0.2% trifluoroacetic acid for 10 minutes and subjected to an additional phosphopeptide enrichment step using High-Select™ Fe-NTA spin columns (Thermo Fisher Scientific, catalog # A32992) to reduce the abundance of non-phosphorylated peptides which may have bound nonspecifically to the IP beads. Phosphopeptides were eluted according to the manufacturer’s instructions, dried by vacuum centrifugation, and resuspended in 5% acetonitrile in 0.1% formic acid.

### Liquid chromatography and mass spectrometry

TMT-labeled phosphopeptides were loaded directly onto a chromatography column using a helium packing device in order to minimize sample loss. Columns were constructed in-house; fused silica capillary with inner diameter 50 μm and outer diameter 375 μm (Molex, catalog # 1068150017) was cut to a length of 25 cm and pulled using a micropipette laser puller to create an integrated emitter tip with 1-2 μm inner diameter, then was packed with 10 cm of 3 μm C18 beads (YMC Co, catalog # AM12S03) and conditioned and tested using a tryptic digest of bovine serum albumin. Liquid chromatography-tandem mass spectrometry was performed using an Agilent (1100-Series) chromatograph coupled to a Thermo Scientific™ Orbitrap Exploris™ 480 mass spectrometer. Peptides were separated using 0.2M acetic acid (solvent A) and 70% acetonitrile in 0.2M acetic acid (solvent B) over the following 140-minute gradient profile: (min:%B) 0:0, 10:11, 115:32, 125:60, 130:100, 133:100, 140:0. Electrospray ionization was carried out at 2.5 kV and at a flow rate of approximately 100 nL/min (200 μL/min through a flow splitter achieving a ∼1:2000 split). The mass spectrometer was operated in data-dependent acquisition mode as follows: in each cycle, a full (MS) scan of precursor ions with m/z between 380-1800 was acquired with a resolution of 120,000 at 200 m/z and AGC target of 3e6, followed by a maximum of 20 MS/MS scans per cycle. For tandem (MS/MS) scans, precursors with charge state between 2-5 were isolated with an isolation width of 0.4 Th and accumulated until the AGC target of 1e6 was reached, or until 247 ms had elapsed. Fragmentation was performed using a normalized collision energy of 33%, and fragment ions were scanned with a resolution of 120,000 at 200 m/z. Each precursor was allowed to be selected for MS/MS four times before being dynamically excluded for 3 minutes.

### Phosphopeptide identification and quantitation

Raw files containing phosphopeptide mass spectra were processed using Proteome Discoverer version 3.0 (Thermo Fisher Scientific) and searched using Mascot version 2.4 (Matrix Science). Spectra were searched against the canonical human proteome (SwissProt reviewed sequences, version 2023_04) with trypsin digestion allowing for up to one missed cleavage, and with a precursor ion mass tolerance of 5 ppm and a fragment ion mass tolerance of 20 mmu. Precursor ions and TMT reporter ions were removed from MS/MS spectra prior to searching using the non-fragment filter node in Proteome Discoverer. Cysteine carbamidomethylation, TMT-labeled lysine, and TMT-labeled peptide N-termini were set as static modifications; phosphorylation of tyrosine, serine, and threonine, and oxidation of methionine were set as dynamic modifications. Phosphorylation site localization was performed using ptmRS within Proteome Discoverer (62). For peptide quantitation, TMT reporter ion intensities were extracted with an integration tolerance of 10 ppm and were isotope-corrected using Proteome Discoverer according to the manufacturer-provided batch-specific isotopic impurities of each TMT channel.

Peptide-spectrum matches (PSMs) were exported from Proteome Discoverer and filtered by match quality (expectation value < 0.05) and phosphosite localization confidence (probability > 0.9 for all phosphosites on the peptide). Any PSM from a spectrum that matched to more than one amino acid sequence with equal expectation value was removed. PSMs with missing values in more than half of the TMT channels were discarded. All remaining PSMs were subjected to replacement of missing TMT reporter ion intensities with a value equivalent to half the intensity of the least-intense fragment ion observed in the corresponding MS/MS spectrum (25). TMT reporter ion intensities were then summed across PSMs sharing a common modified peptide sequence. To correct for technical variation in the total amount of peptide labeled in each TMT channel, a loading control factor was calculated for each channel by subjecting crude peptide from the phosphotyrosine IP supernatant (approximately 50 ng) to mass spectrometry according to the procedure outlined above. Phosphopeptides without at least one phosphorylated tyrosine were discarded. Phosphopeptides derived from the same source protein(s) with shared phosphorylation site pattern (for example, two phosphopeptides covering the same phosphosite but with different tryptic cleavage pattern) were aggregated to minimize phosphosite redundancy in the resulting data matrix.

### Statistical analysis and visualization

All computational and statistical analyses were performed with preexisting functions from the SciPy (60) and seaborn (63) python packages as described in the Materials and Methods section. For the kinase activity analysis shown in Figure 4B, protein-protein interaction data was downloaded from BioGRID version 4.4 (37). pY sites with unique mapping to a single protein and with significant difference between GTL and GSL (greater than 1.5-fold and *P* < 0.05 by two-sided *t*-test with multiple hypothesis correction by the Benjamini-Hochberg procedure) were considered for kinase activity inference. The schematics depicted in Figures 2A, S1A, and S1D were created using BioRender and can be accessed at https://BioRender.com/t8r352v.

## Data availability

All dose-response cell viability data, drug combination cell viability data, and processed pY proteomics data will be made available as supplementary material. The raw and searched mass spectrometry pY proteomics data will be made available on the ProteomeXchange Consortium via the PRIDE partner repository.

## Acknowledgements

The authors would like to thank the following individuals for helpful discussions and critical feedback throughout this work: Douglas Lauffenburger, Michael Hemann, Ryan Sullivan, Jason Conage-Pough, Benjamin Neel, Sabrina Spencer, Uwe Rix, Juhyeon Son, Ryuhjin Ahn, and Tigist Tamir. This work was supported by NIH grant U54 CA283114 and the MIT Center for Precision Cancer Medicine. C.T.F. was funded by a graduate fellowship from the Ludwig Center at MIT.

## Author contributions

C.T.F. and F.M.W. designed research; C.T.F. performed research; C.T.F. and F.M.W. analyzed data; C.T.F. wrote the original draft; and C.T.F. and F.M.W. wrote the paper.

---

**Figure S1.**
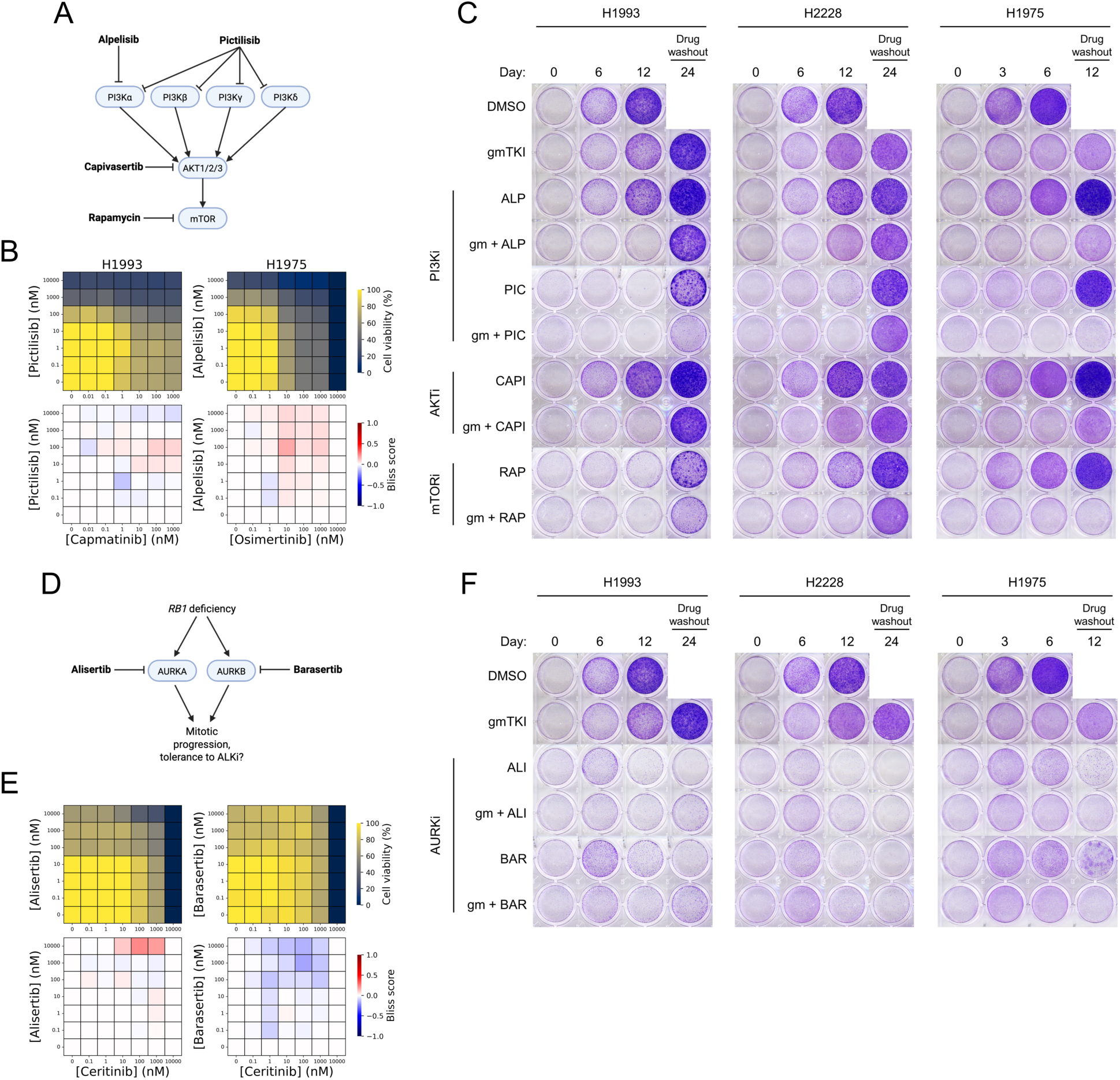
A subset of gmTKI-tolerant lines are sensitive to combined PI3K/AKT/mTOR pathway inhibition. (A) Pathway schematic depicting the targets of inhibitors used in the experiments shown in (B) and (C). (B) Dose-resolved cell viability measurements (top) and corresponding Bliss synergy matrices (bottom) from checkerboard assays pairing gmTKI with PI3K inhibitors in H1993 (left) and H1975 (right) cells. Treatment duration was 72 hours. Cell viability values were averaged over six biological replicates and normalized to DMSO control. (C) Time-resolved clonogenic assays of GTLs treated with gmTKI, PI3K/AKT/mTOR pathway inhibitors, or combinations thereof. All drugs were delivered at a dose of 1 μM. Timing of the experiment was compressed for H1975 due to its significantly faster growth which caused poor cell adhesion under certain drug treatments after 10-12 days. (D) Pathway schematic depicting the targets of AURK inhibitors used in the experiments shown in (E) and (F). (E) Dose-resolved cell viability measurements (top) and corresponding Bliss synergy matrices (bottom) from checkerboard assays pairing gmTKI (ceritinib) with AURK inhibitors in H2228 cells. Treatment duration was 72 hours. Cell viability values were averaged over six biological replicates and normalized to DMSO control. (F) Time-resolved clonogenic assays of GTLs treated with gmTKI, AURK inhibitors, or combinations thereof. All drugs were delivered at a dose of 1 μM.

**Figure S2.**
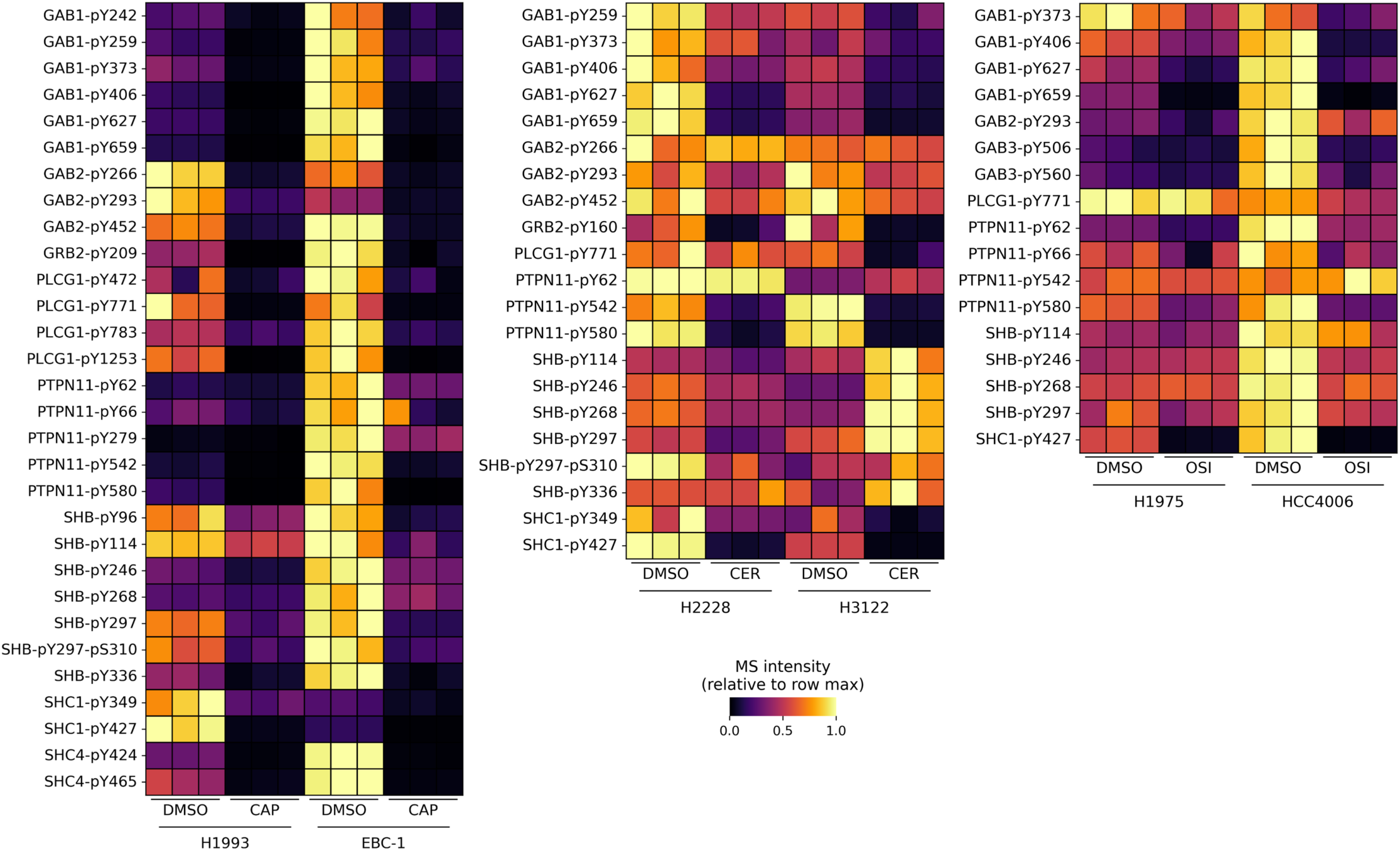
GTLs and GSLs undergo similar remodeling of their downstream driver TKs under gmTKI treatment. Heatmaps depicting the relative abundance of pY sites on adapter proteins and other downstream signaling effectors in (left) MET-amplified cell lines, (middle) ALK-rearranged cell lines, and (right) EGFR-mutant cell lines under DMSO or gmTKI.

**Figure S3.**
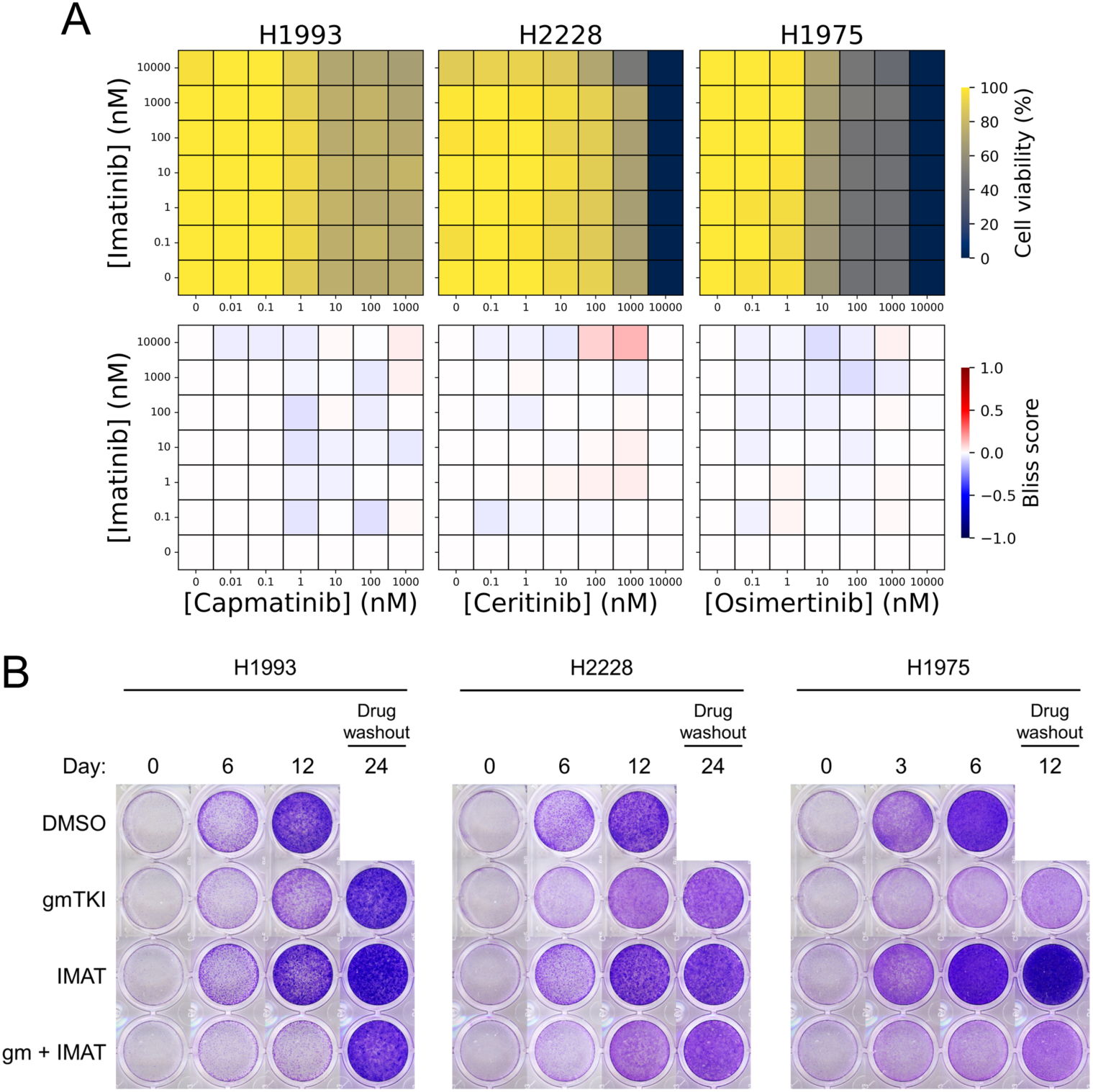
ABL inhibition without SFK inhibition shows modest effect in lines with elevated ABL signaling. (A) Dose-resolved cell viability measurements (top) and corresponding Bliss synergy matrices (bottom) from checkerboard assays pairing gmTKI with imatinib, an ABL1/2 inhibitor with selectivity over SFKs. Treatment duration was 72 hours. Cell viability values were averaged over six biological replicates and normalized to DMSO control. (B) Time-resolved clonogenic assays of GTLs treated with gmTKI, imatinib, or the combination. All drugs were delivered at a dose of 1 μM except imatinib, which was delivered at 10 μM due to its higher IC_50_ for ABL1/2. Timing of the experiment was compressed for H1975 due to its significantly faster growth which caused poor cell adhesion under certain drug treatments after 10-12 days.

